# MethodsJ2: A Software Tool to Improve Microscopy Methods Reporting

**DOI:** 10.1101/2021.06.23.449674

**Authors:** Joel Ryan, Thomas Pengo, Alex Rigano, Paula Montero Llopis, Michelle S. Itano, Lisa Cameron, Guillermo Marqués, Caterina Strambio-De-Castillia, Mark A. Sanders, Claire M. Brown

## Abstract

Proper reporting of metadata is essential to reproduce microscopy experiments, interpret results and share images. Experimental scientists can report details about sample preparation and imaging conditions while imaging scientists have the expertise required to collect and report the image acquisition, hardware and software metadata information. *MethodsJ2* is an *ImageJ/Fiji* based software tool that gathers metadata and automatically generates text for the methods section of publications.

## ARTICLE

Optical microscopy is used in nearly all fields of research spanning from life and health sciences to many areas of physical sciences and engineering. The lack of reproducibility in science is a widespread problem which leads to significant challenges for researchers, slows scientific progress and wastes valuable resources.^1–3^ To improve reproducibility there needs to be detailed reporting of both research resources^4^ and experimental methods^2^. Progress has been made with tools to promote and enable antibody validation^1,5,6^, cell line authentication^7–11^ and identify reagents and tools/services through the Research Resource ID (RRID) (https://scicrunch.org/resources). RRIDs are used to report antibodies, model organisms, cell lines and plasmids in addition to custom software, databases and services (e.g. core facilities such as imaging platforms). There are not many tools for experimental methods reporting and it remains a difficult challenge to solve.

A lack of methods reporting is a widespread problem in microscopy where many articles contain no information or lack basic details about how images were collected^12^. Analysis of 240 research articles published in 8 mainstream journals containing ~3,000 figures, of which more than half included images, revealed that only 17% of the publications passed a test for minimal information required to reproduce the experiment^12^. The problem is compounded by the sheer number and variety of microscope modalities, options and associated components, such as the light source, optics and detectors. In addition, advances in microscopy have automated the process to a level that has distanced the researcher from the technical parameters. Finally, while researchers are focused on scientific questions under study and have extensive expertise with their model systems (e.g. sample preparation, imaging conditions) they typically do not have an in-depth background in microscopy. As a result, it is difficult for experimental scientists (i.e. microscope users) to be aware of what information needs to be reported to enable proper evaluation and reproduction of their work.

Essentially, to properly evaluate microscopy data and ensure it is reproducible, information about sample preparation (e.g. tissue, cell type, dye), experimental conditions (e.g. temperature, live, fixed), microscope hardware (e.g. objective lens, filters, camera), image acquisition settings (e.g. exposure time, pixel size), quality control metrics (e.g. light source stability, resolution) and image analysis parameters (e.g. segmentation, background correction) used to generate the images and any quantitative results is required. This information is called “metadata” and is defined as “a set of data that describes and gives information about other data”. Researchers involved in the 4D Nucleome initiative^13^ and Bioimaging North America (BINA) (https://www.bioimagingna.org/) have developed extensive community driven *Microscopy Metadata* specifications^14,15^. These specifications build on a previous Open Microscopy Environment (OME) model^16^ and include an in-depth community driven *Microscopy Metadata* model for light microscopy termed “4DN-BINA-OME”^14,17^. The model scales with experimental design, instrument complexity and the degree to which image processing and quantitative image analysis is required for interpreting the results. This ensures that only essential information required to reproduce each type of imaging experimental results is included to minimize the burden on experimental scientists to collect, annotate and report metadata. The umbrella term for metadata information is *Image Metadata* that is then classified into different subtypes including *Experimental and Sample Metadata*, *Microscopy Metadata* and *Analysis Metadata*. *Microscopy Metadata* includes hardware specifications, image acquisition settings and image structure (pixel size, number of pixels, planes, colours and dimensions)^18^.

To help solve the complex problem of methods reporting papers have been published on establishing minimal and accurate microscopy information guidelines^17,19,20^, information for reporting image processing^21^, what can go wrong if detailed metadata is not reported^22^ and the importance of measuring and reporting microscope quality control ^23^. Improving awareness and education around *Image Metadata* and how it is essential for reproducible microscopy experiments is important. However, to really tackle the problem and make a significant impact, it is vital to have straightforward readily accessible tools for implementation by experimental scientists.

This manuscript presents *MethodsJ2*, an extensible open source microscopy methods reporting software tool that runs in *ImageJ/Fiji* and builds on our recently published work (*MethodsJ*)^12^. Fundamentally, *MethodsJ2* captures *Image Metadata* from multiple sources, consolidates it and automatically generates a detailed methods text for publication. Integration with *ImageJ/Fiji* was specifically chosen to make it broadly available and particularly straightforward to experimental scientists to incorporate it into their imaging workflows.

Once an image is open in *ImageJ/Fiji*, *MethodsJ2* automatically gathers metadata from the image using OME BioFormats (e.g. camera exposure time, pixel size, magnification). It then captures *Microscopy Metadata* from a Microscope.JSON file generated using *Micro-Meta App*^15,24^. *Micro-Meta App* is a software tool that guides imaging scientists or microscope custodians step-by-step in the collection of *Microscopy Metadata* associated with a specific microscope that is consistent with community standards^14^ and stores it in a Microscope.JSON file. This file only needs to be generated once and updated if microscope hardware is upgraded or replaced. Normally a specialist (e.g. imaging scientist) will use *Micro-Meta App* to set up configurations for each microscope they manage and will provide experimental scientists with a Microscope.JSON. Next, *MethodsJ2* guides the user to manually enter specific *Experimental and Sample Metadata* (e.g. cell type, dyes, live or fixed samples). The researcher is then prompted and guided step-by-step through all collected metadata for validation and modification if needed. Imaging scientists can automatically integrate acknowledgements text for their imaging facility (including a RRID) into the *MethodsJ2* script so it is included in the manuscript. This will considerably improve publication tracking to monitor and demonstrate facility impact on science. Finally, the methods text is generated and can be reviewed and finalized for publication.

Another complementary software tool to facilitate *Microscopy Metadata* reporting is *OMERO.mde*^25^. This tool focuses on consistent handling of *Image Metadata* ahead of data publication as specified by shared community *Microscopy Metadata* specifications^14,16^ and according to the FAIR principles^26,27^. It can be used for the early development and maturation of image metadata extension specifications to maximize flexibility and customization while at the same time allow for testing and validation before incorporation in community-accepted standards.

### Detailed *MethodsJ2* workflow (Figure 1)

**Figure 1:**
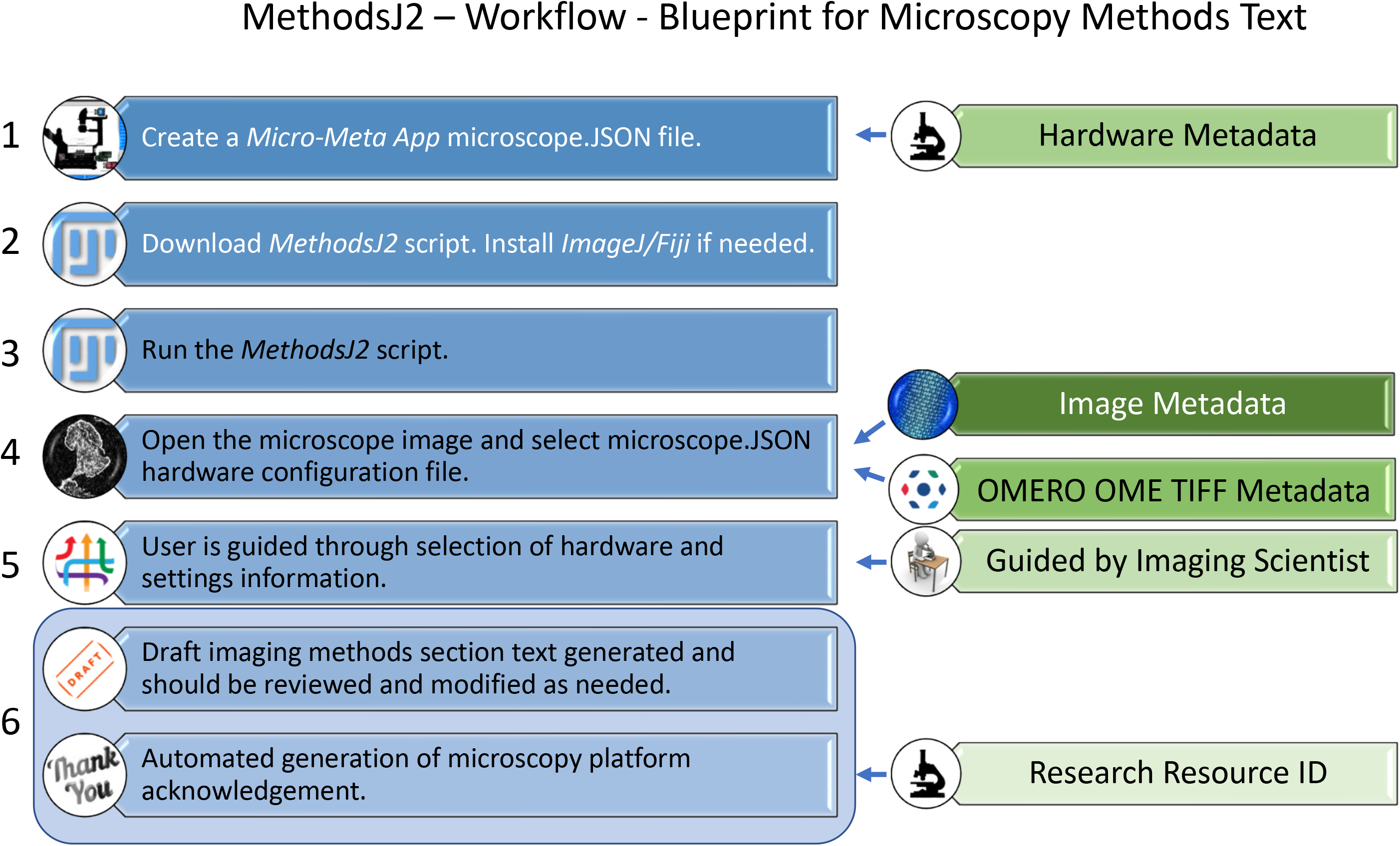
*MethodsJ2* Workflow Overview. Steps required to automatically generate microscopy methods text. Image metadata is collected from the manufacturer metadata in the image file using the OME TIFF tools. Hardware metadata is collected from a *Micro-Meta App* Microscope.JSON file. It is recommended to have an experienced microscopist or imaging scientist guide researchers through the methods text generation and validation process.

#### Note

An in-depth workflow with software screenshots is available as supplemental material.

#### Step 1

Use *Micro-Meta App* to create and save a Microscope.JSON file including all of the *Hardware Metadata*. This is a time-consuming process but is only done once, typically by a microscope expert. <u>**Note:**</u> When creating the Microscope.JSON file it is important to give each component a detailed name as this will be used to populate text information in *MethodsJ2*. For example, put “63x/1.4 NA Plan-Apochromatic oil immersion” not “63x”.

#### Step 2

Download the *MethodsJ2* script, an example Microscope.JSON file and an example image from GitHub (https://github.com/ABIF-McGill/MethodsJ2). If needed download and install *ImageJ/Fiji* (https://fiji.sc/).

#### Step 3

Drag the *MethodsJ2* script file and drop it on the *ImageJ/Fiji* toolbar. It will automatically open in the Script Editor and from there press “*Run*”.

#### Step 4

*MethodsJ2* will prompt the user to open an image to use to generate the microscopy methods text. The *Image Metadata* is automatically extracted by *MethodsJ2* using Bio-Formats and microscopy manufacturer proprietary image formats. Then the user is prompted to select a *Microscope.JSON* file for the corresponding microscope use to generate the image.

#### Step 5

*MethodsJ2* will prompt the user for sample information, then guide the user step-by-step to select and validate the image and hardware settings used to generate the selected image based on metadata extracted from the image and Microscope.JSON file.

#### Critical Step

Have an experienced microscope user or imaging scientist from a microscopy platform guide the researcher through the experimental, software and hardware settings information for validation. Any missing information can be manually added based on published community guidelines.

#### Step 6

Following validation click “*OK*”. Draft text is automatically generated and appears in a popup window, copied to the clipboard, and can pasted into a manuscript. <u>**Note:**</u> It is important to review the draft text to ensure that it is accurate and make minor adjustments for grammar.

Comprehensive methods reporting is essential for image analysis workflows^28,29^, data analytics such as statistical analysis^30^, reusability of imaging data in public archives^31–33^ and when applied to emerging artificial intelligence-based image analysis methods^34,35^. Continued progress along the path of rigor and reproducibility are essential for high quality research data for the expanse of researchers using microscopy to make new discoveries and broadly share image data. There is a shared responsibility to continue to make improvements to ensure quality and reproducibility. Experimental scientists must use due diligence to understand the fundamentals of the technologies their research relies on and work with imaging scientists to ensure the required metadata is collected and reported. Imaging scientists need to support and educate experimental scientists, so they understand what metadata needs to be reported and why. Microscope manufacturers need to integrate, automate and report *Microscope Metadata*. Scientific publishers and reviewers have a duty to promote community-based guidelines^16,25^ including the 4DN-BINA-OME microscopy metadata model^14^ and ensure microscope data that is published meets a minimum standard. It is in the best interest of funding agencies to uphold high quality and reproducible microscopy image data and make certain that detailed *Microscopy Metadata* is available when image data is shared to harness the maximal amount of information and discovery for the broader research community and the public.

*MethodsJ2* and two companion software tools, *Micro-Meta App24* and *OMERO.mde*^25^, advance rigor and reproducibility in microscopy. The implementation and evolution of all three of these tools will help all stakeholders ensure *Microscopy Metadata* is documented and reported until a more fully integrated solution is found. These tools immediately promote transparency and reproducibility in microscopy. However, there are still many challenges. A great deal of hardware and software information is recorded by proprietary microscope manufacturer image acquisition software. However, this information is not typically available to microscope users because the software code is not open source. In addition, when it is recorded in image files the information is not provided in standard formats. This problem is compounded when images are saved and opened with a third-party software and any metadata - however limited - is often lost^14^. In fact, images collected with the default settings from many different microscope manufacturers provide very limited *Microscope Metadata*^14^ and rely on microscope users to accurately input the information. Microscope manufacturers need to work with the global community and organizations such as the newly created group Quality Assessment and Reproducibility for Instruments & Images in Light Microscopy (QUAREP-LiMi)^36^ to automate metadata collection, ensure it conforms to community standards^14,16,25^ and make it readily available.

As with any software tool, work is ongoing. Future development will: 1) integrate the *Micro-Meta App* software metadata *Settings.JSON* file into *MethodsJ2*. 2) Adapt *Micro-Meta App* to generate methods text directly so researchers can use their platform of choice. 3) Adapt *MethodsJ2* to automatically extract extended experimental metadata from *OMERO.mde*. and 4) Expand all three tools to include both advanced microscopy modalities (e.g. confocal, super-resolution, intravital) and calibration and performance metadata (e.g. resolution and illumination stability metrics) on the basis of the 4DN-BINA-OME Microscopy Metadata community driven specifications.

## Supporting information

Supplemental instructions

## ACKNOWLEDGEMENTS

We thank our microscopy core facility staff and users of the University Imaging Centers of the University of Minnesota (RRID:SCR_020997); McGill University Advanced BioImaging Facility (ABIF) (RRID:SCR_017697), the UNC Neuroscience Microscopy Core (RRID:SCR_019060) (supported, in part, NIH-NINDS Neuroscience Center Support Grant P30 NS045892, NIH-NICHD Intellectual and Developmental Disabilities Research Center Support Grant U54 HD079124) and the MicRoN (Microscopy Resources on the North Quad) Core at Harvard Medical School. Chan Zuckerberg Initiative DAF, an advised fund of Silicon Valley Community Foundation supports C.M.B. (Grant# 2020-225398), C.S. D. C. (Grant# 2019-198155 (5022)) and M.S.I. (Grant# 2019- 198107).

---

**Supplemental Figure 1:**
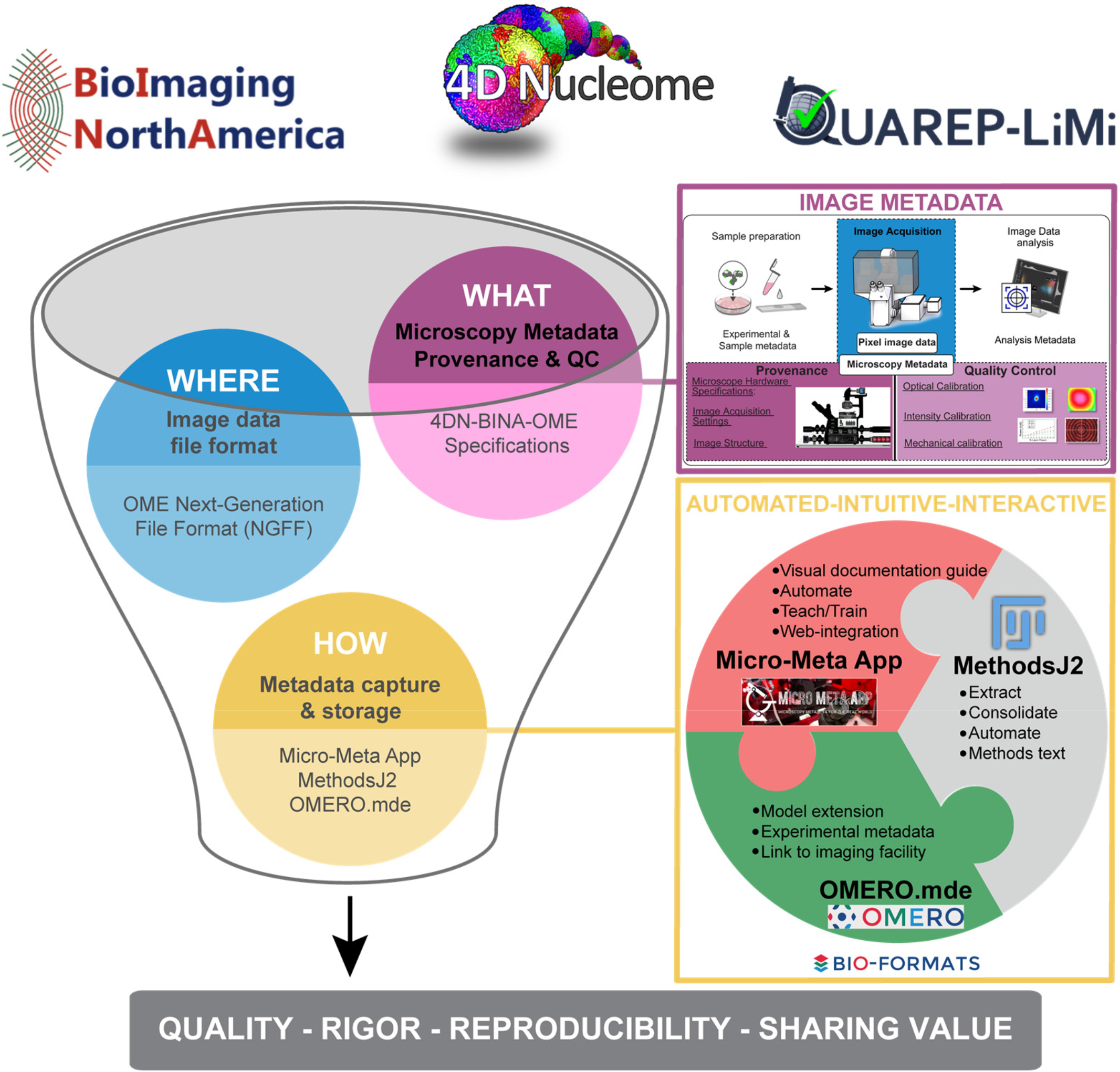
Quality, rigor, reproducibility and sharing value for imaging experiments require the definition of community-driven Microscopy Metadata specifications and the adoption of easy-to-use metadata collection tools to facilitate the documentation and quality control tasks for experimental scientists. The establishment of FAIR (Wilkinson et al., 2016), community-driven Microscopy Image Data Standards implies parallel development on three interrelated fronts: 1) Next-Generation File Formats (NGFF) <u>where</u> the ever-increasing scale and complexity of image data and metadata would be contained for exchange (Moore et al., 2021); *blue bubble*). 2) Community-driven specifications for <u>what</u> ‘data provenance’ information (microscope hardware specifications, image acquisition settings and image structure metadata) and quality control metrics are essential for rigor, reproducibility, and reuse and should therefore be captured in Microscopy Metadata (*magenta bubble*). 3) Shared rules for <u>how</u> the (ideally) automated capture, representation and storage of Microscopy Metadata should be implemented in practice (*yellow bubble*). Micro-Meta App, MethodsJ2 and OMERO.mde are three highly interoperable and complementary tools that function to train users on the importance of documentation and quality control, facilitate metadata extraction, collection and storage, and automatically write Methods sections. The different tools are based on different software platforms in order to appeal to the broadest community including microscope builders, imaging scientists working in core facilities and experimental scientists. The concept is to bring the tools to software platforms people are already using and lower the bar to enable broad uptake.

## Notes

https://github.com/ABIF-McGill/MethodsJ2

### Competing Interest Statement

The authors have declared no competing interest.

