## Supplemental instructions for "MethodsJ2: A Software Tool to Improve Microscopy Methods Reporting"

### MethodsJ2 step-by-step overview

https://github.com/ABIF-McGill/MethodsJ2

### MethodsJ2 - Workflow

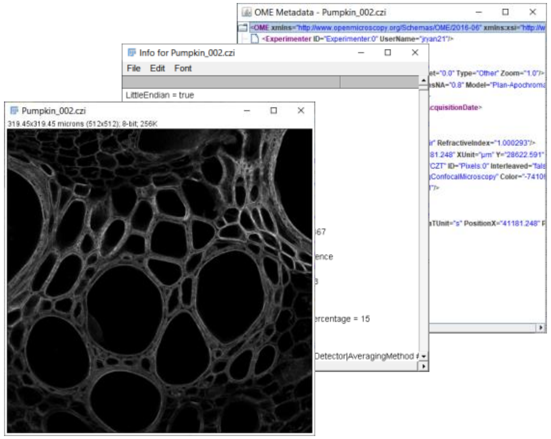


Image, metadata,
OME metadata

MethodsJ2
structure file for
dialog boxes and
text generation

**MethodsJ2**

Python script
running in Fiji

User input,
guided by core
facility staff

Generates materials and
methods section for
imaging experiments,
based on community
guidelines

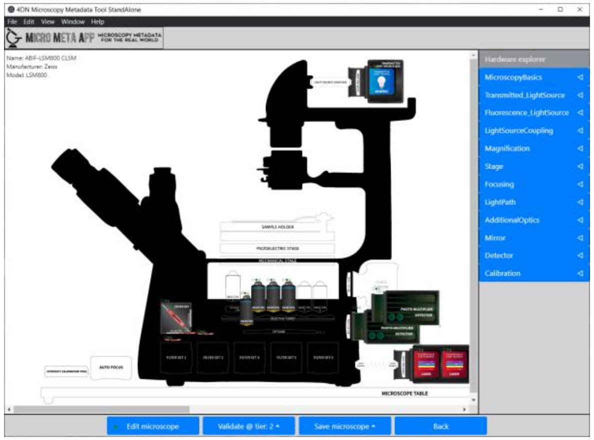


Micro-Meta App microscope
hardware specifications file

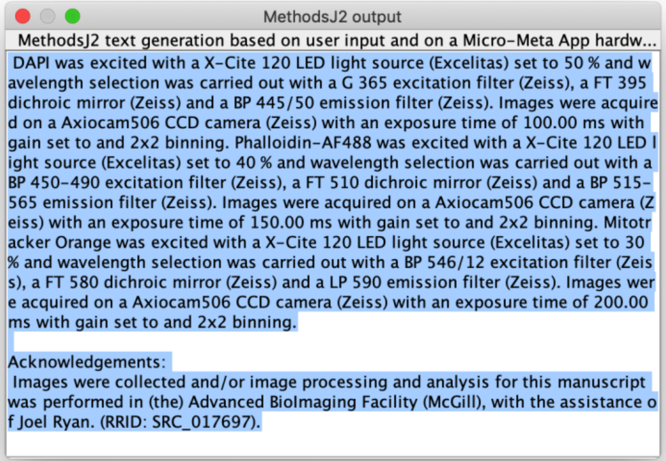

### MethodsJ2 - run

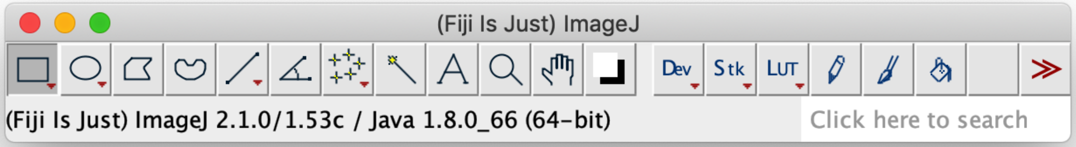


Drag and drop MethodsJ2.py on the main Fiji toolbar. Alternatively, click File > New > Script, then in the Script Editor, Click File > Open, and select MethodsJ2.py

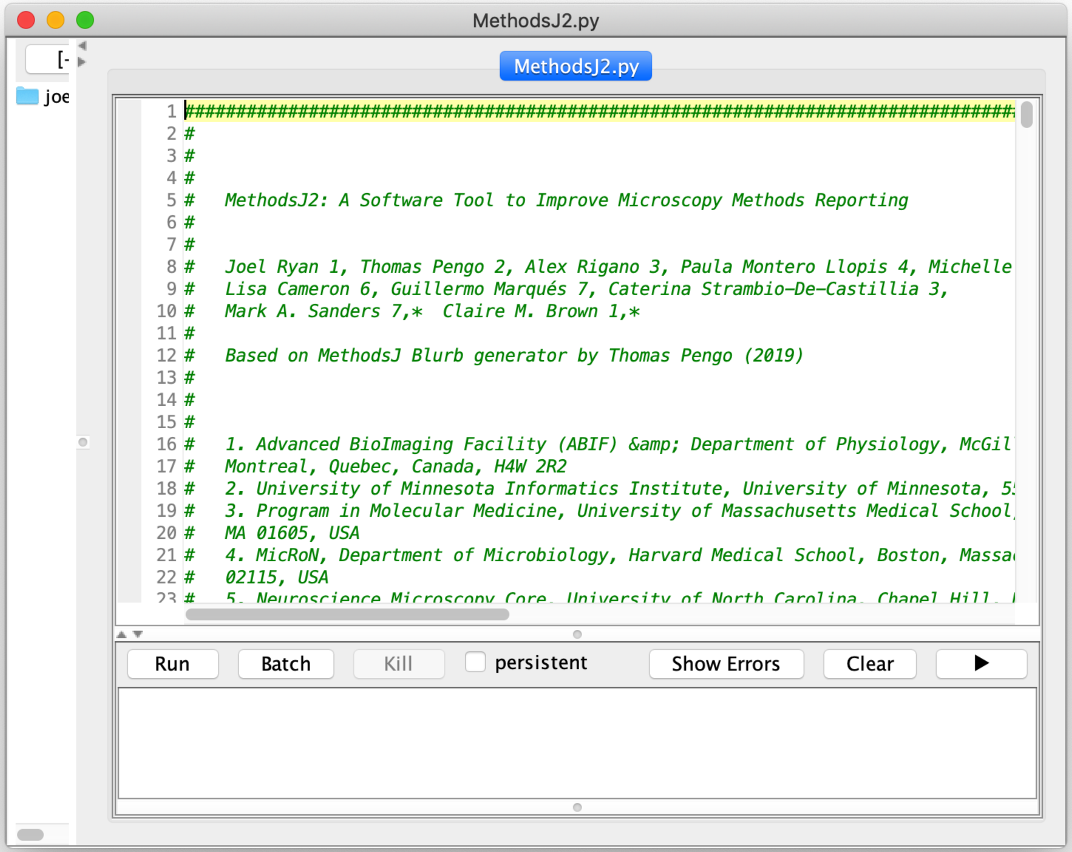


Check language: click Language and select Python

Once the script is loaded is ready, click Run

It may take a few seconds to start.

### MethodsJ2 – select image

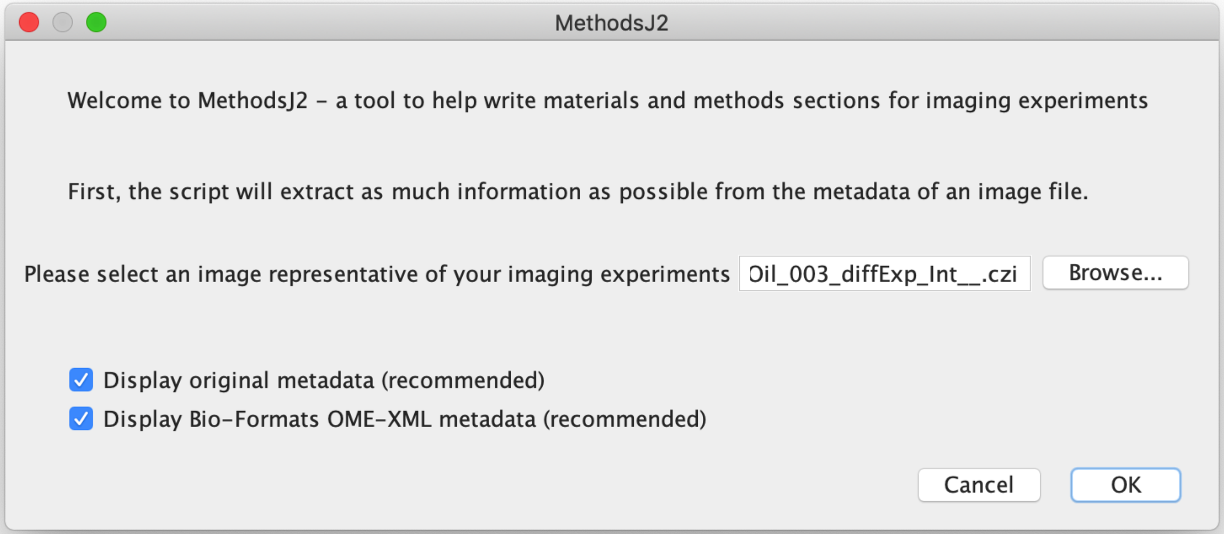


Select an image to load and to source metadata.

Click on Browse and navigate to the image, or drag and drop in image file into the text input field

Optional: display metadata windows (useful for filling out dialog boxes later)

### MethodsJ2 – select image

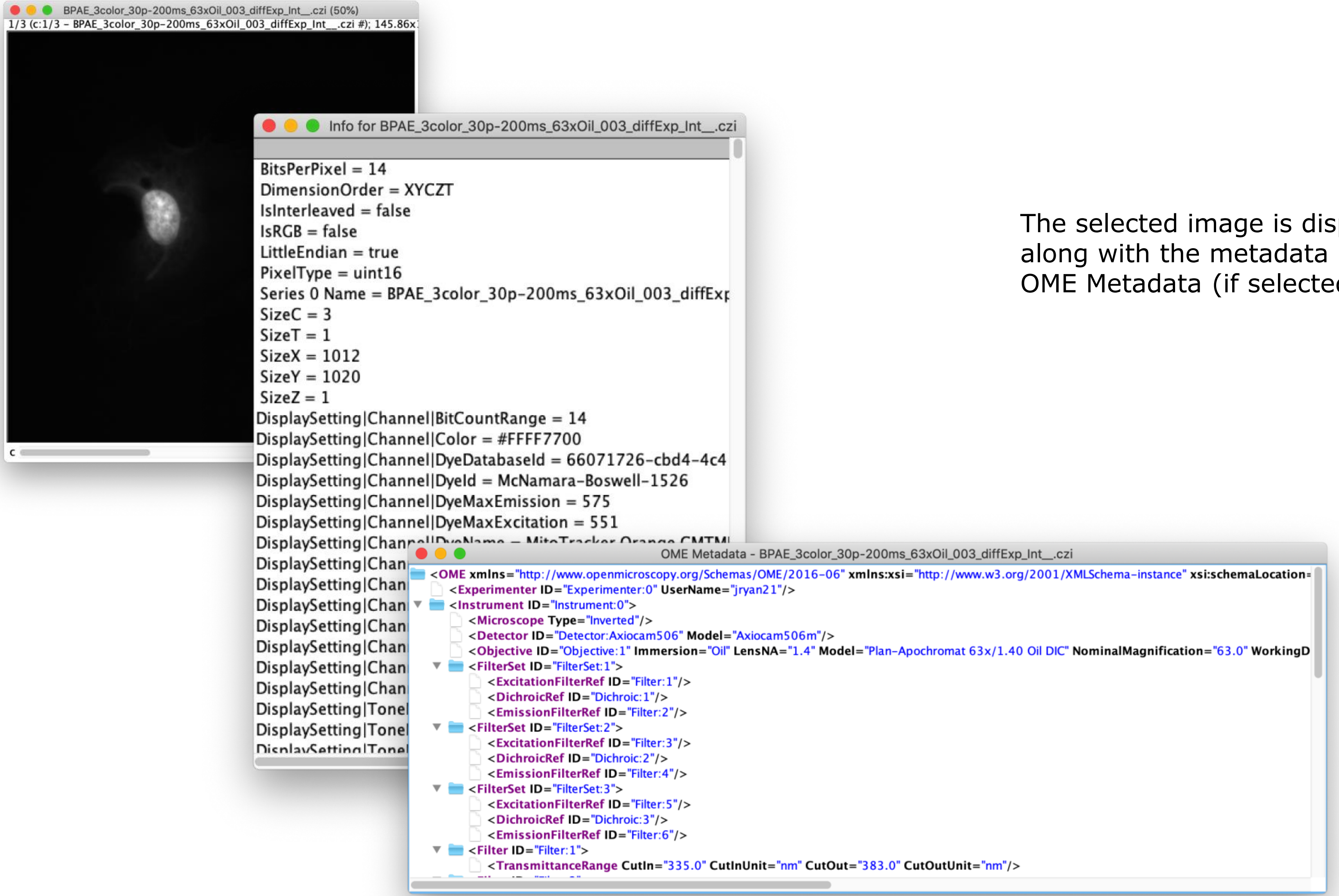


The selected image is displayed, along with the metadata and OME Metadata (if selected)

### MethodsJ2 – Sample Preparation

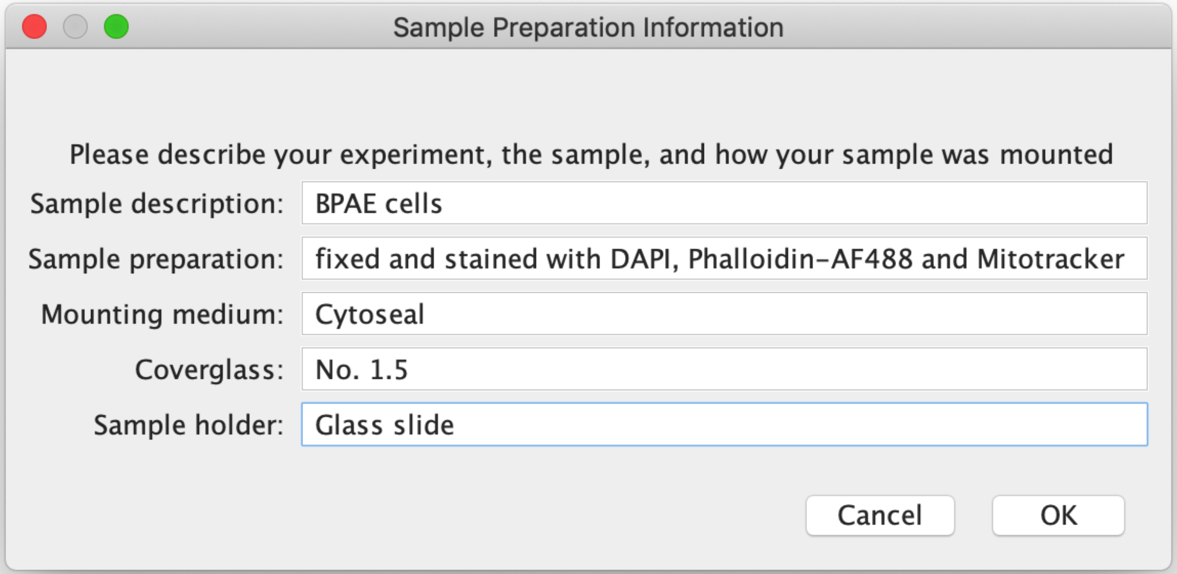


Please provide information about the sample, and how it was prepared for imaging

Given the variety of samples and preparations, no text is generated for sample description. It is more of a reminder for users to provide complete sample information.

### MethodsJ2 – Image dimensions

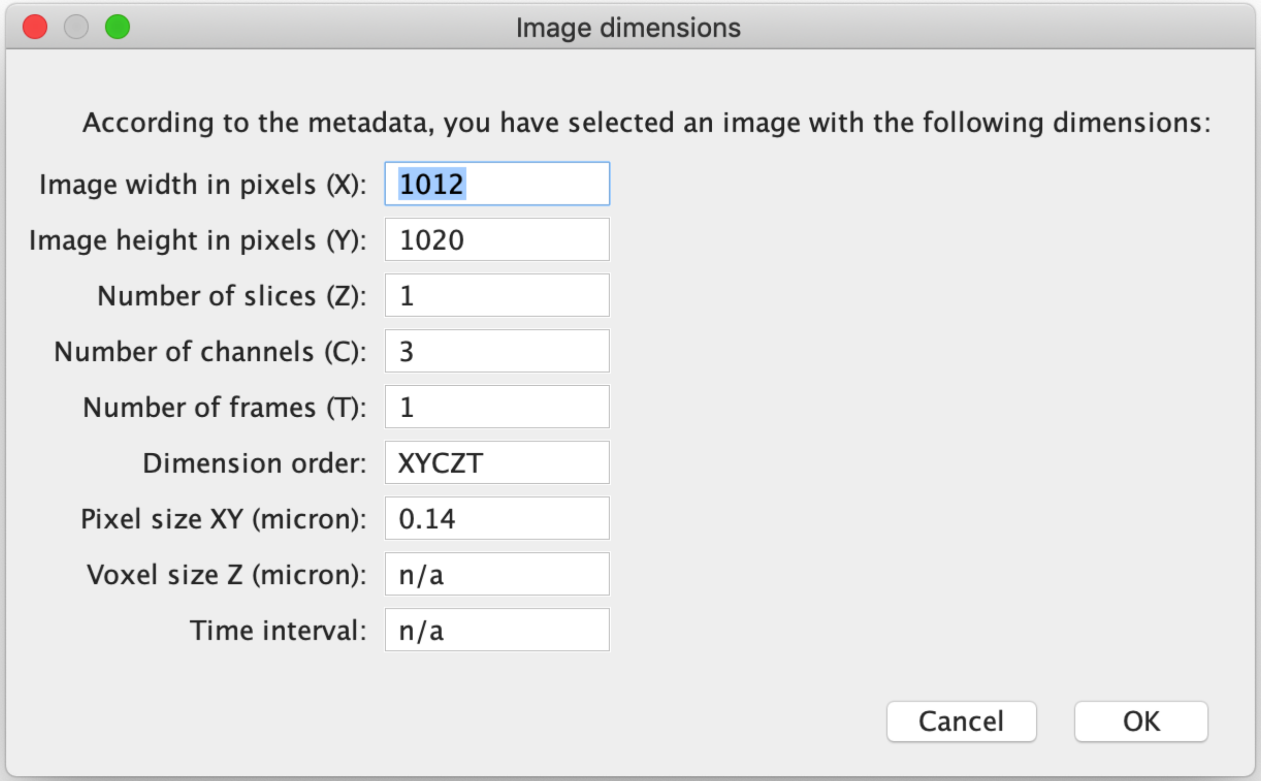


Please verify image dimensions. Values are sourced from the image metadata

### MethodsJ2 – select Microscope.json file

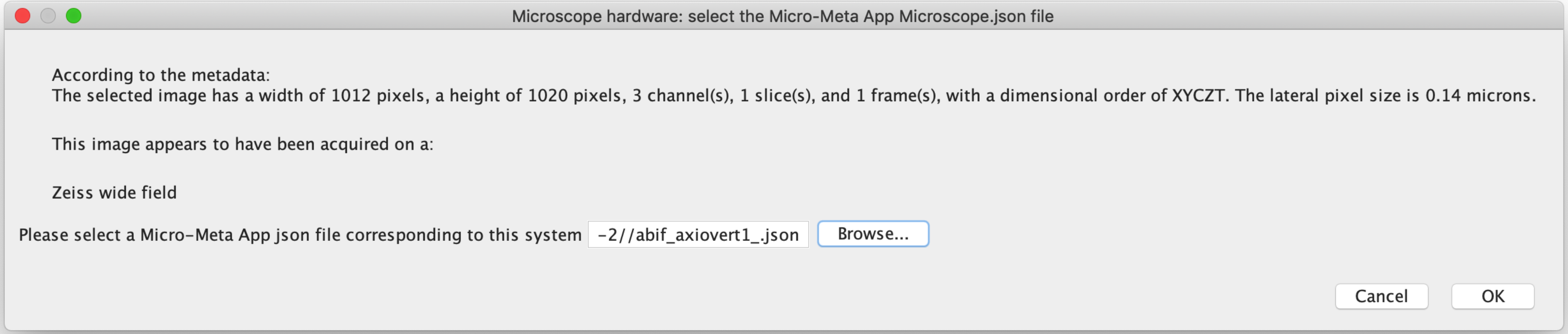


Choose Micro-Meta App hardware specifications file for the microscope used to acquire the selected image

### MethodsJ2 – choose descriptor and software

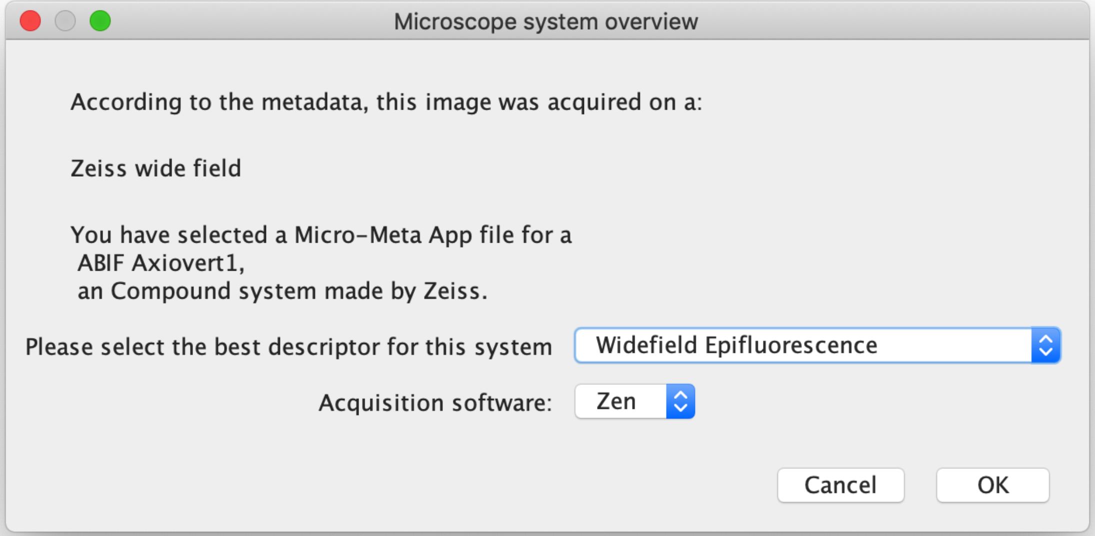


Please select the best descriptor for the selected microscope, as well as the acquisition software.

### MethodsJ2 – select objective

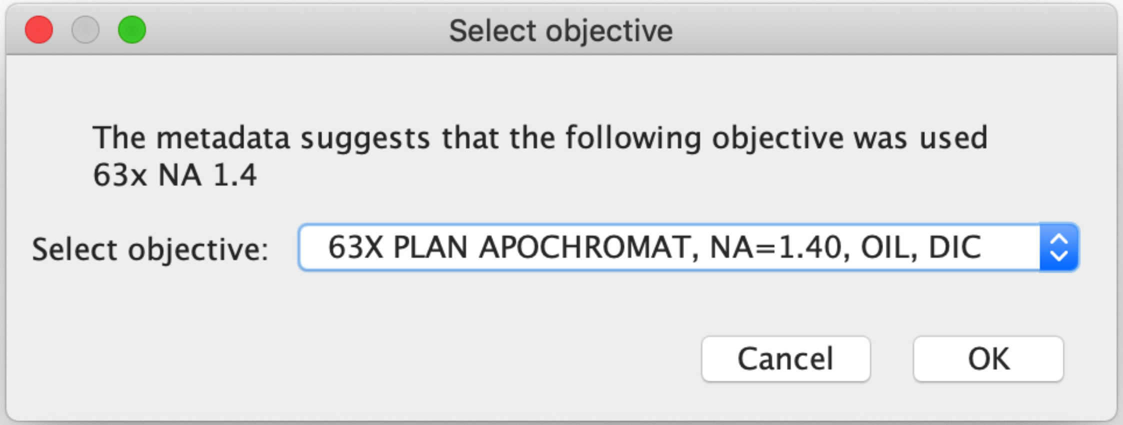


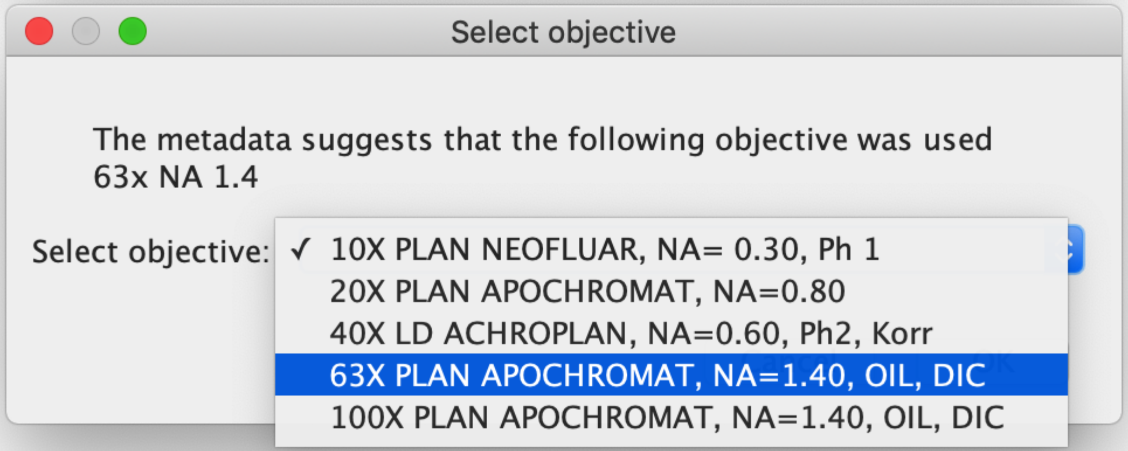


Select the objective used for this experiment.

A suggestion is made based on the metadata, and the list of objectives to choose from is sourced from the microscope configurations file.

The drop-down menu is populated from objectives available in the Micro-Meta app hardware specifications file.

### MethodsJ2 – Channel acquisition settings

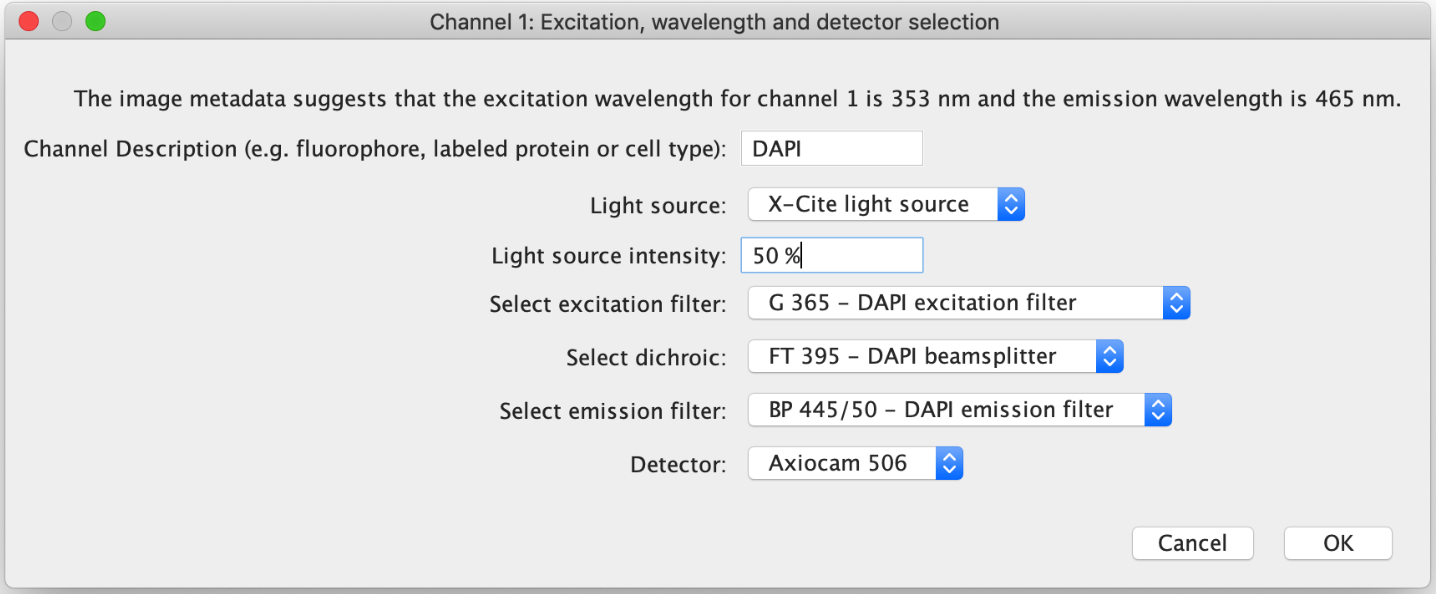


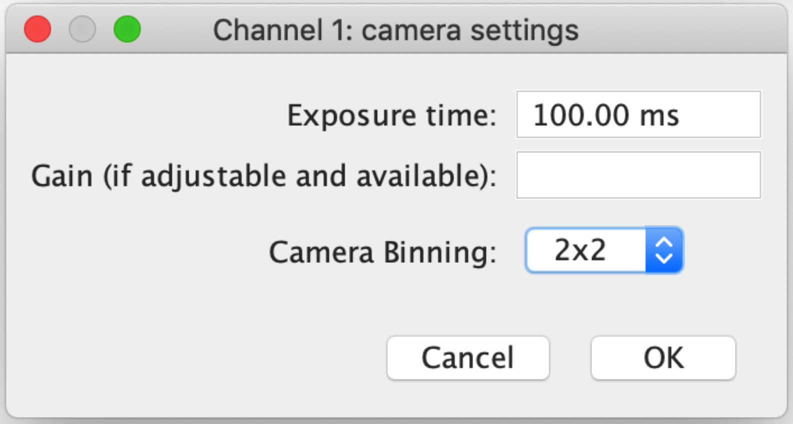


Please fill in information for the first channel

Options in drop-down menus are sourced from the microscope configuration file.

Detector settings are based on whether a camera or point detector is selected

### MethodsJ2 – Channel acquisition settings loop

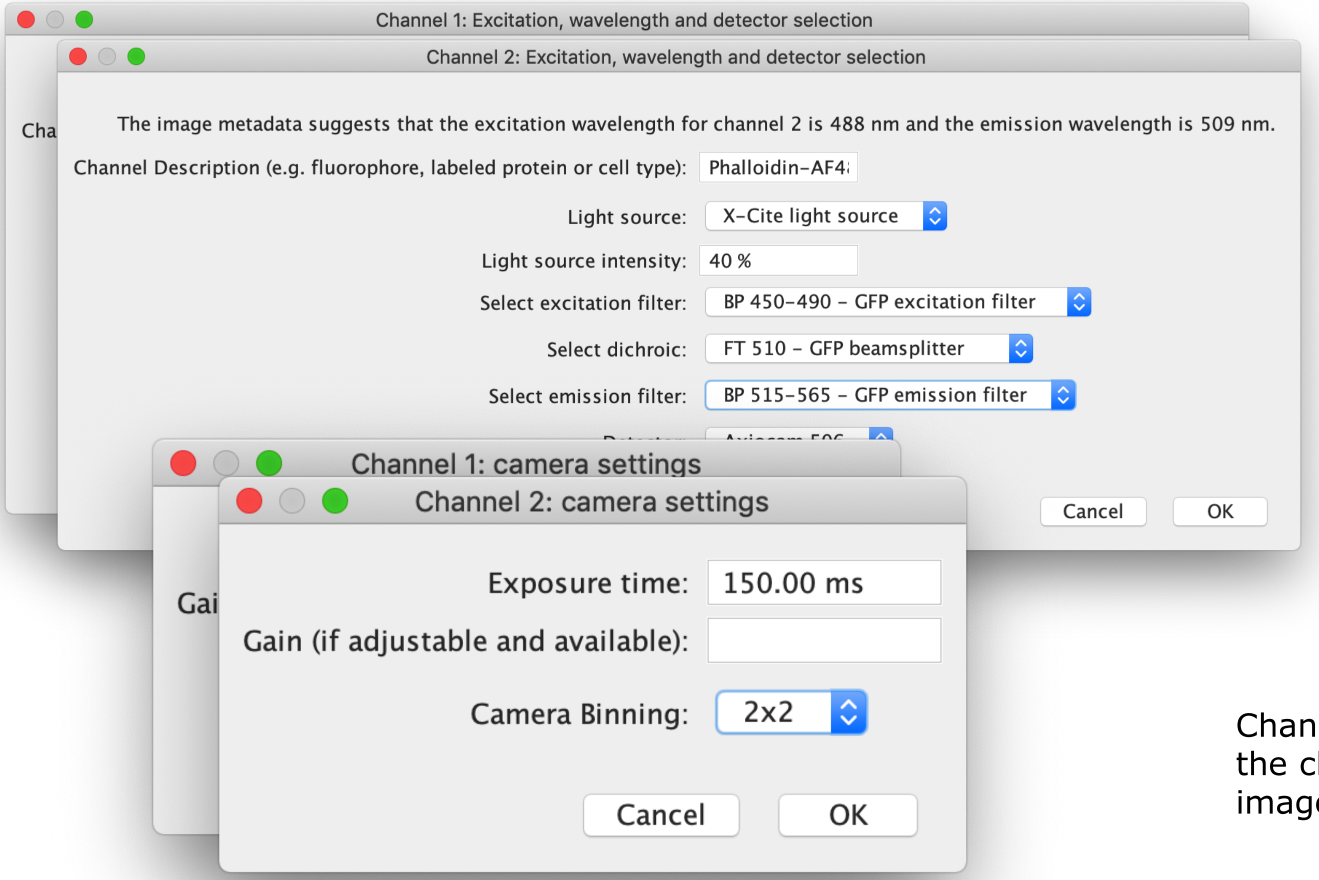


Channel menus will loop through the channels in the selected image.

### MethodsJ2 – select optional devices

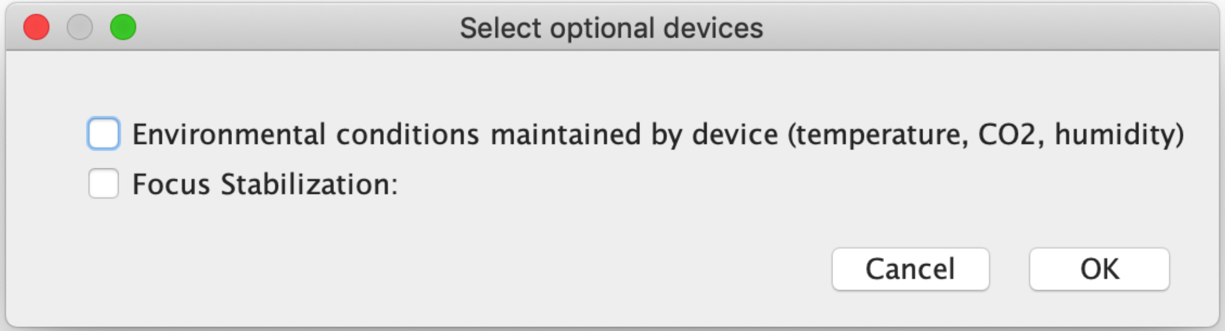


Choose whether optional devices from the microscope hardware specifications file were used for the selected image.

### MethodsJ2 – Sample text for acknowledgement

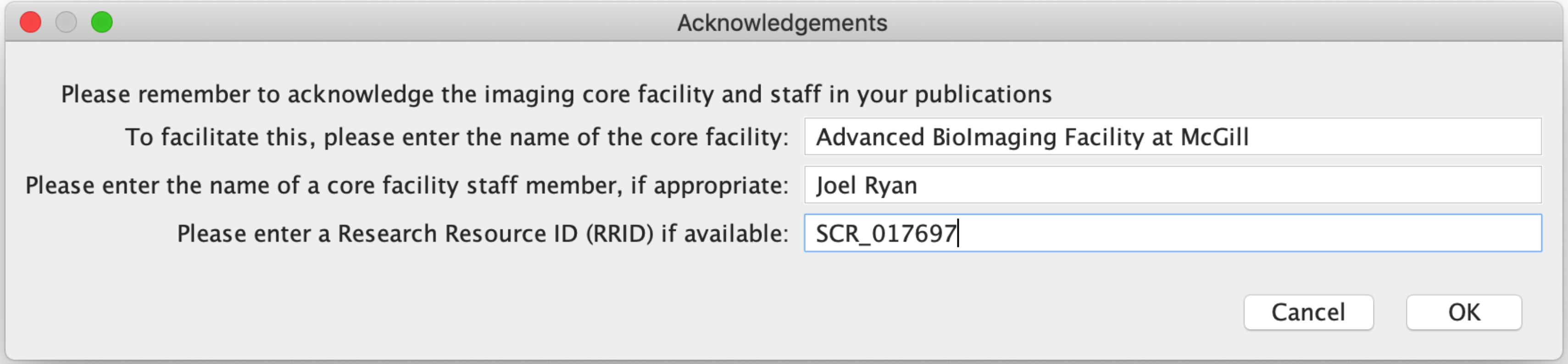


Please enter the name of the core facility or laboratory which manages the microscope used for the acquisition of the selected image, as well as any imaging scientist who was helpful in the imaging experiment, and if available a Research Resource ID

### MethodsJ2 – output

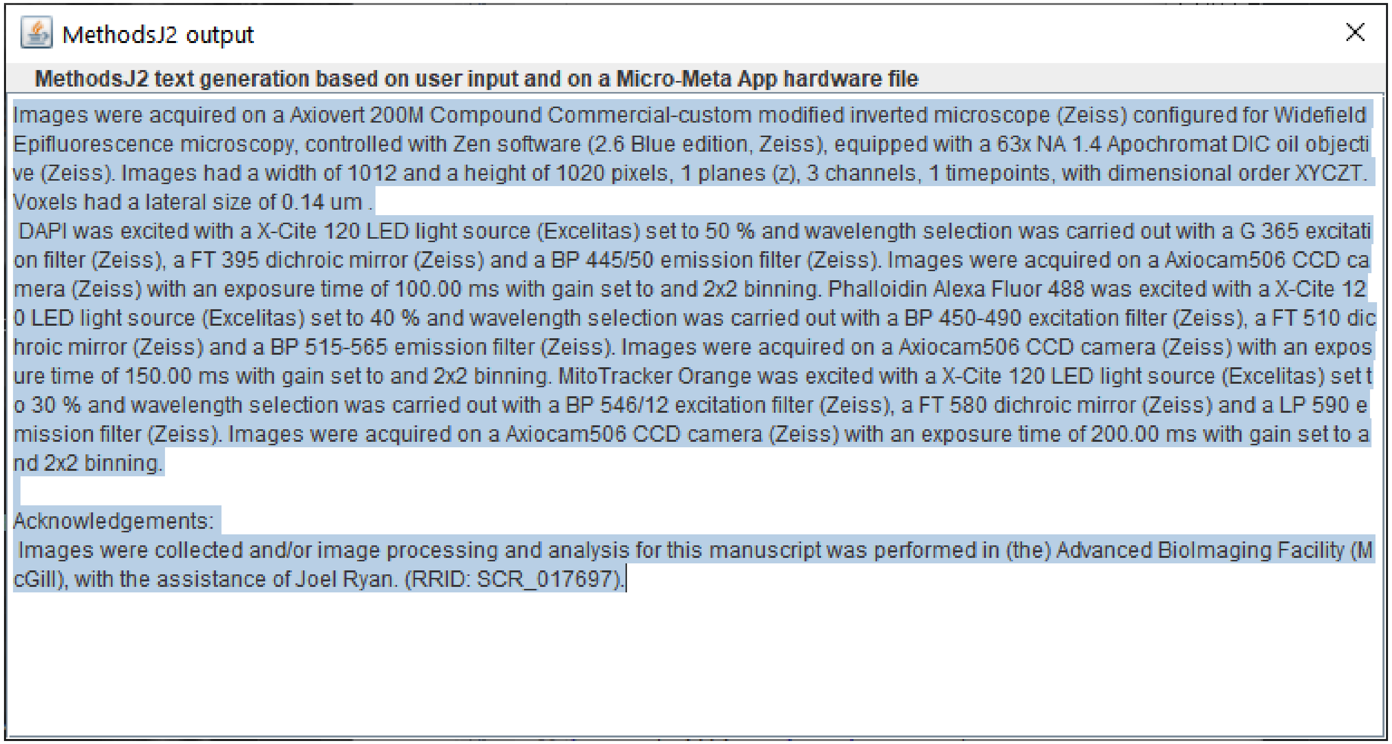


A draft of an experimental section text is displayed in a popup window and copied to the clipboard, to be pasted into a manuscript for revision.

### MethodsJ2 – extensibility


```json
 {
    "Dialog_Box": "Channel Settings",
    "category": "general",
    "Setting": "Light source intensity: ",
    "Add_to_same_row": 0,
    "CheckHardwareJSON": 0,
    "Dialog_Type": "addStringField",
    "blurb": "set to %s"

},
{
    "Dialog_Box": "Channel Settings",
    "category": "general",
    "Dialog_Type": "addChoice",
    "Setting": "Select excitation filter: ",
    "Add_to_same_row": 0,
    "CheckHardwareJSON": 1,
    "Schema_ID": "ExcitationFilter.json",
    "attributes": [
        "Model",
        "Manufacturer"
    ],
    "blurb": "and wavelength selection was carried out with a %s excitation filter (%s), "
},
```


Dialog boxes, drop-down menus, text generation can be added or modified by core facility staff, by modifying a MJ2 structure file and storing it locally or online (e.g. on Github)

https://raw.githubusercontent.com/ABIF-McGill/MethodsJ2/main/MJ2_structure_files/MJ2_structure_file_001.json
